# Fitness dependence preserves selection for recombination across diverse mixed mating systems

**DOI:** 10.1101/2020.09.29.318238

**Authors:** Sviatoslav Rybnikov, Daniel B. Weissman, Sariel Hübner, Abraham B. Korol

## Abstract

Meiotic recombination and the factors affecting its rate and fate in nature have inspired many theoretical studies in evolutionary biology. Classical theoretical models have inferred that non-zero recombination can be favoured under a rather restricted parameter range. Thus, the ubiquity of recombination in nature remains an open question. However, these models assumed constant (uniform) recombination with an equal rate across all individuals within the population. Models of fitness-dependent recombination, with the rate varying among genotypes according to their fitness have shown that such a strategy can often be favoured over the best constant recombination. Here we use simulations to show that across a range of mating systems with varying frequencies of selfing and clonality, fitness-dependent recombination is often favoured even when any non-zero constant recombination is disfavoured. This recombination-protecting effect of fitness dependence is strongest under intermediate rates of selfing or high rates of clonality.

## 1. Introduction

Meiotic recombination can generate new potentially beneficial allele combinations but simultaneously destroy the existing ones that have already proved to be successful. Given this well-recognized evolutionary ambiguity of recombination, its evolution has long remained a challenging question. Classical models for the evolution of recombination usually assume infinite panmictic populations, a small number of loci, and constant recombination rate. Despite the limitations of these models due to analytical constraints, they were able to provide highly valuable theoretical results. According to these models, non-zero recombination can be favoured under relatively stringent constrains on selection parameters (first and foremost, selection intensity and epistasis) that are not easily found in nature [1]. This finding appeared in a contradiction with the ubiquity of recombination in nature, which prompted further theoretical research. An apparent solution was to consider more realistic assumptions including finite and/or structured populations, and multi-locus genetic systems [2], which do indeed expand the parameter space compatible with non-zero recombination rate [3–5]. Alternatively, the theory should take into consideration the essential features of recombination, such as the sensitivity of recombination rate to external (ecological stressor) and/or internal (genotype fitness) conditions. Indeed, models with such condition-dependent recombination suggest that it can be more advantageous than constant recombination under various scenarios [6–12].

In a previous study, we considered an infinite population of diploid individuals subjected to purifying epistatic selection [13]. In such setup, fitness-dependent recombination often appeared to be favoured *even when any positive constant recombination was disfavoured*. Here, we extend the model to explore the conditions favouring fitness-dependent recombination also under other mating strategies. We begin with a marginal situation of complete outcrossing and gradually move towards complete self-fertilization or clonality. We first identify selection regimes disfavouring constant recombination and then assess in which proportion of such ‘recombination-unfriendly’ conditions recombination can still be favoured if its rate varies among genotypes according to their fitness. The pair of recombination strategies in question are compared in terms of modifier approach [14], i.e., based on allele dynamics at a selectively neutral locus only affecting recombination rates between the selected loci. However, we also evaluate the effect of fitness-dependent recombination on several important population-level characteristics including the time needed to reach the equilibrium state (mutation–selection balance), the average fitness at equilibrium, and the variance of fitness at equilibrium.

## 2. Models and methods

### (a) Genetic system and selection regime

All individuals across all models are diploids. Each individual bears three to five bi-allelic loci affecting fitness (hereafter ‘selected loci’). The wild-type allele *A* ensures fitness of 1, while the mutant allele *a* decreases it to 1–*SH* and 1–*S* in the hetero- and homozygous states, respectively. Thus, the parameters *S* and *H* denote the deleterious effect of mutations and their dominance, respectively. If a genotype bears mutant alleles at several selected loci, its fitness decreases more pronouncedly than expected upon independent effects (synergistic, or negative epistasis). The epistasis arises in three types of interactions: additive-by-additive (*E_a×a_*), additive-by-dominance (*E_a×d_*), and dominance-by-dominance (*E_d×d_*). The number of the corresponding epistatic interactions in a given genotype can be calculated as follows [5]:

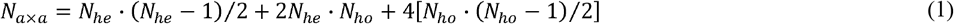

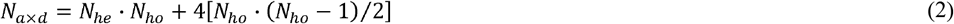

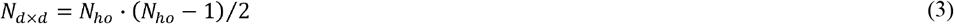

where *N_he_* and *N_ho_* stand for the number of selected loci bearing mutations in the hetero- and homozygous state, respectively. The overall fitness of the genotype is [5]:

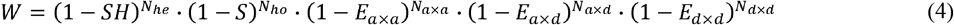

Thus, selection regimes can be formalized as points in five-dimensional space, with coordinates representing five selection parameters (the deleterious effect of mutations, their dominance, and the three epistatic components).

### (b) Life cycle and mating systems

We consider infinite populations with non-overlapping generations. During the maturation from zygotes to adults, genotypes are subject to mutations and selection. The adults then produce zygotes of the next generation. Let 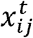 be the frequency of zygote *ij* (i.e., zygote comprised of haplotypes *i* and *j*) in generation *t*. Due to mutations this frequency changes, as follows:

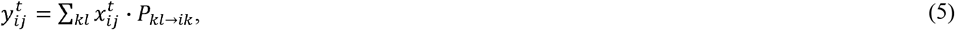

where *P_kl→ij_* is the probability to obtain genotype *ij* from genotype *kl*. For each selected locus, we assume mutations from allele *A* to allele *a* with the frequency of 10^-4^ and no backward mutations. Then, the frequency further changes due to selection, with viabilities defined by *W_ij_*:

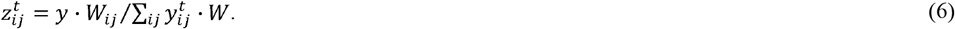

The adults reproduce via outcrossing, selfing, or clonal (asexual) propagation. Upon these three cases, the frequency of zygote *ij* in generation *t* + 1 can be calculated as follows:

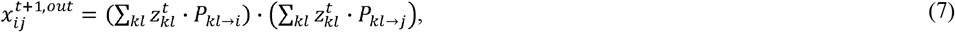

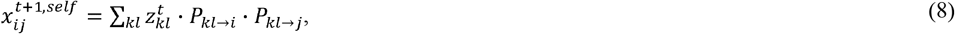

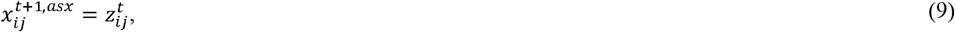

where *P_kl→i_* is the probability to obtain gamete of haplotype *i* from the adult of genotype *kl*. Upon a mixed mating system, the frequency is:

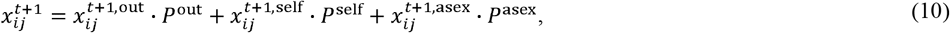

where *P*^out^, *P*^self^, and *P*^asex^ are the probabilities of the adult to reproduce via outcrossing, selfing, and clonal propagation, respectively (*P*^out^ + *P*^self^ + *P*^asex^ = 1).

Such formalization allows modelling various mixed mating systems. Here we consider the two main axes: from complete outcrossing to either complete selfing or almost complete clonality (up to 99%, since complete clonality leaves no place for the evolution of recombination). The probabilities of different forms of reproduction are treated as parameters, not functions of genotype at a specific modifier locus.

### (c) Recombination strategies and their comparison

Recombination rates between the selected loci are controlled by a selectively neutral modifier locus. For simplicity, recombination rates in all pairs of adjacent selected loci are assumed equal, and the crossover events do not interfere. The modifier locus is unlinked to the selected loci. Modifier alleles confer different recombination strategies: constant (uniform) recombination rate across all genotypes, or fitness-dependent recombination with the varying rate among genotypes according to their fitness.

To compare any two recombination strategies, we allowed the selected system to evolve from the wildtype state (no mutant alleles at any selected locus) to mutation-selection balance (diagnosed when frequencies of all selected alleles changed less than by 10^-12^ per generation). After this ‘burn-in’ period of several hundred generations, we introduced the modifier allele in question in Hardy-Weinberg equilibrium and linkage equilibrium with the selected genotypes and let the population evolve for 10,000 generations. We say that a strategy is favoured if its allele increased in frequency from rarity (0.05) to abundance (0.95). More complicated outcomes may include stable polymorphism, in which the two competing alleles both increase in frequency from rarity, and bistability, in which they both increase in frequency from abundance.

For each selection regime, we first compared different constant recombination strategies and chose only those regimes where the best constant recombination rate was zero. For such ‘recombination-disfavouring’ regimes, we tested whether fitness-dependent recombination can nevertheless be favoured. We modelled a fitness-dependent recombination rate *r* varying from 0 (in the fittest genotype) to *r*_max_ (in the least fit genotype):

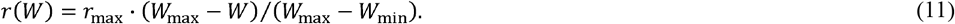

where *W*_min_ and *W*_max_ stand for the minimum and maximum fitness among genotypes heterozygous for at least two selected loci (‘recombination-responsive’ genotypes), since recombination in other genotypes causes no changes in the population structure of the next generation. We considered three different values of *r*_max_: 0.05, 0.1 and 0.2. All plots show results for *r*_max_ = 0.1; the lower (0.05) and higher (0.2) values shift the results slightly, with no effect on the overall patterns.

## 3. Results

First, we examined 10,000 five-dimensional vectors each representing a random selection regime with coordinates distributed uniformly from 0 to 1. This set was applied for each of the considered mating strategies. However, with this set, many biologically interesting parameters combinations remain underrepresented. Therefore, we examined an additional set of 1,000 random parameter combinations with values of the deleterious effect of mutations and three epistatic components distributed uniformly from 0 to 0.1. This parameter area is thus covered by a 1000-times higher density of points compared with the first set.

For each mating strategy, we first identified selection regimes disfavouring non-zero constant recombination (section a) and then tested whether fitness-dependent recombination is favoured under such recombination-disfavoured conditions (section b). Using binary logistic regression, we link the evolutionary advantage or disadvantage of each of the recombination strategies to the corresponding values of selection parameters. Next, we quantified the frequency with which fitness-dependence strategy “rescues” recombination in situations where constant recombination is disfavoured (section c). Finally, we consider the effects of fitness-dependent recombination on several important population-level characteristics (section d).

### (a) Selection for/against non-zero constant recombination

Under *complete outcrossing*, the evolutionary advantage or disadvantage of non-zero constant recombination was determined mostly by three selection parameters: deleterious effect of mutations (*S*), their dominance (*H*) and additive-by-additive epistasis (*E_a×a_*). The two other epistatic components, additive-by-dominance (*E_a×d_*) and dominance-by-dominance epistasis (*E_d×d_*), were negligible. Even such a simple explanatory model demonstrated very high classification accuracy with circa 92% of cases predicted correctly (Table S1A). To further explore interactions between selection parameters, we applied stepwise regression for their different combinations. The resulting model included additive-by-additive epistasis, selection against heterozygotes *SH*, and the triple product *SHE_a×a_*. As seen from the clear-cut hyperbolic patterns in Fig. 1A (top row), the relation between *S* and *H* was inverse; at that, non-zero constant recombination was favoured under relatively high (above a certain threshold) values of *SH*. The interaction between *SH* and *E_a×a_* appeared more complex (Fig. 1A, bottom row) which presumably constrained the stepwise procedures from neglecting the triple product *SHE_d×d_*. Assuming the above-mentioned parameter combinations further increased the classification accuracy, up to ~99%.

**Figure 1:**
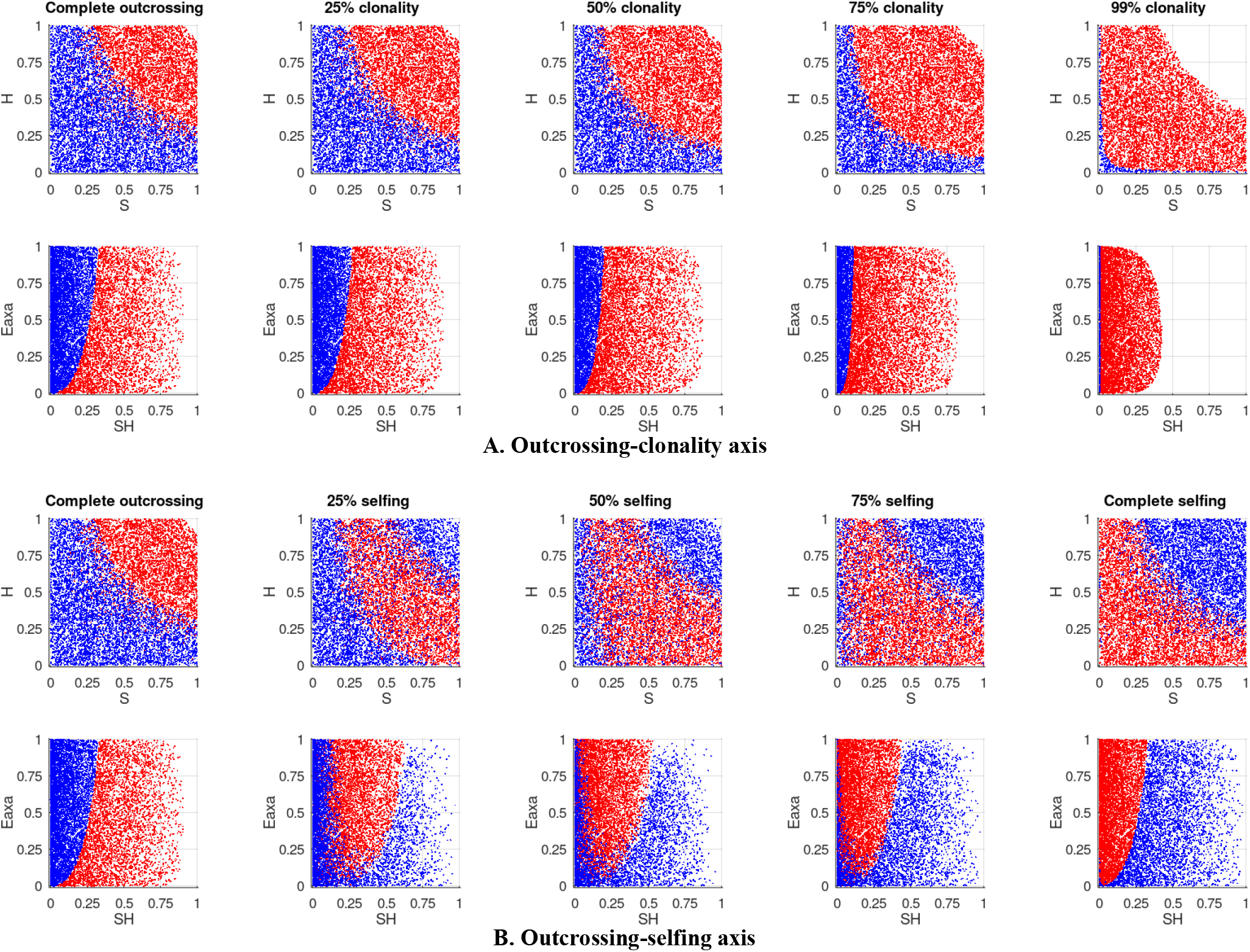
The effects of the main selection parameters on the evolutionary advantage/disadvantage of *non-zero constant recombination* under different mating systems along the outcrossing–clonality (A) and outcrossing–selfing (B) axes. The two rows of scatter plots within each axis represent the most informative coordinate planes: *S–H* (the top row) and *SH–E_a×a_* (the bottom row). The coloured points in the scatter plots represent selection regimes that favour constant recombination (red) and disfavour it (blue). Here and in following figures, all data shown are simulated with three selected loci.

The above-described interactions between the selection parameters remained consistent along the *outcrossing–clonality axis*, although additive-by-additive epistasis gradually lost its influence (Table S1A). During this transition, non-zero constant recombination was favoured under lower values of *SH* (Fig. 1A), which decreased the proportion of ‘recombination-disfavouring’ regimes (Fig. 2A, blue curve). Interestingly, the same three selection parameters that determined the evolutionary advantage or disadvantage of non-zero constant recombination under complete outcrossing (*S, H, E_a×a_*) remained the most influential also under *complete selfing* (Table S1B). However, the direction of their effect inverted: non-zero constant recombination was favoured under relatively low values of *SH* (Fig. 1B, top row). As a result, along the *outcrossing-selfing axis*, the area of ‘recombination-disfavouring’ regimes gradually moved from low to high values of *SH* (Fig. 1B) while their proportion did not change drastically (Fig. 2B. blue curve). During this transition, the evolutionary advantage of non-zero constant recombination was determined by a non-linear and sometimes even non-monotonous interplay between the selection parameters. Notably, under intermediate rates of selfing, additive-by-dominance epistasis became more influential. Consequently, classification accuracy under intermediate rates of selfing was reduced to circa 72% (Table S1B).

**Figure 2:**
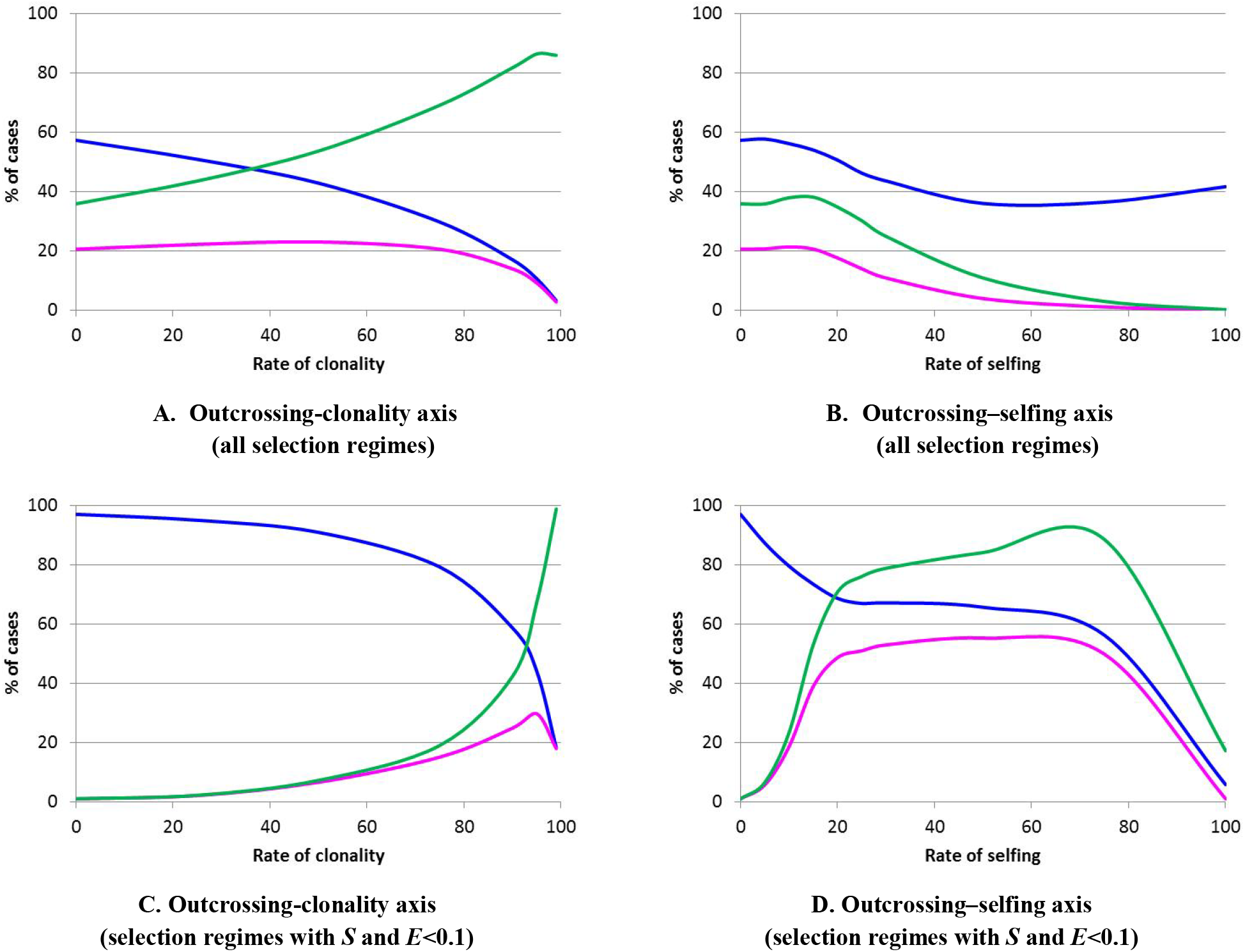
The recombination-rescuing potential of fitness-dependent recombination, under different mating systems along the outcrossing–clonality (A and C) and outcrossing–selfing (B and D) axes. The coloured curves show the proportion of selection regimes disfavouring any positive constant recombination (blue); the proportion of selection regimes disfavouring any positive constant recombination but favouring fitness-dependent recombination (magenta); and the relative proportion of regimes favouring fitness-dependent recombination among the set of regimes disfavouring positive constant recombination (green). The top subplots (A and B) show averages over all selection regimes while the bottom ones (C and D) are restricted to more biologically realistic regimes, with *S* and *E*<0.1.

In the more biologically realistic part of parameter space, non-zero constant recombination was almost always disfavoured under complete outcrossing. In contrast, under high rates of clonality or complete selfing, non-zero constant recombination was favoured in most cases (Fig. 1). As a result, the overall tendencies for the area of realistic selection appeared slightly different, with more pronounced decreasing slopes across both outcrossing–clonality and outcrossing–selfing axes (compare blue curves in Figs. 2A and 2B versus 2C and 2D, respectively).

The proportion of cases with polymorphism or bistability strongly depended on the difference in recombination rate of the two competing strategies. For the pair *r*=0 and *r*=0.05, polymorphism was a rare outcome (<0.4%), observed under low-to-intermediate rates of selfing (5–30%) both over the full parameter space and in the area of realistic selection. Situations with bistability were also rare (<0.4%) over the whole parameter space but quite frequent (up to 14%) in the area of realistic selection.

### (b) Selection for/against fitness-dependent recombination

Our main objective was to explore the parameter space in which fitness dependence can rescue recombination from ‘extinction’. Thus, after identifying the parameter values in which positive constant recombination is disfavoured, we simulated a competition between fitness-dependent recombination and zero recombination. Over the full range of the selection parameters, we found that selection for/against fitness-dependent recombination under *complete outcrossing* was determined by additive-by-additive epistasis *E_a×a_*, selection against heterozygotes *SH*, and the triple product *SHE_a×a_* – similarly to selection for/against non-zero constant recombination. To be favoured, fitness-dependent recombination required *SH* to be above a certain threshold (Fig. 3A). The magnitude of recombination plasticity was the fourth most influential parameter, hence the stronger the fitness dependence the more likely that recombination is favoured.

**Figure 3:**
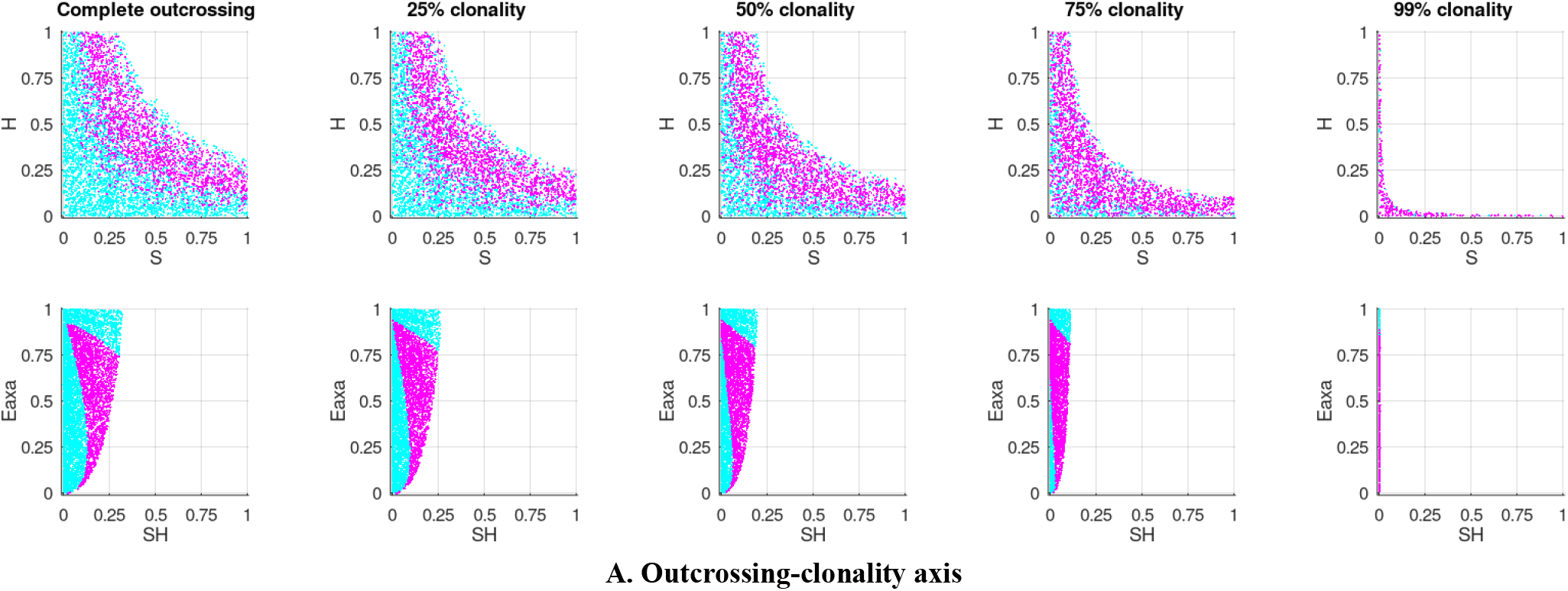
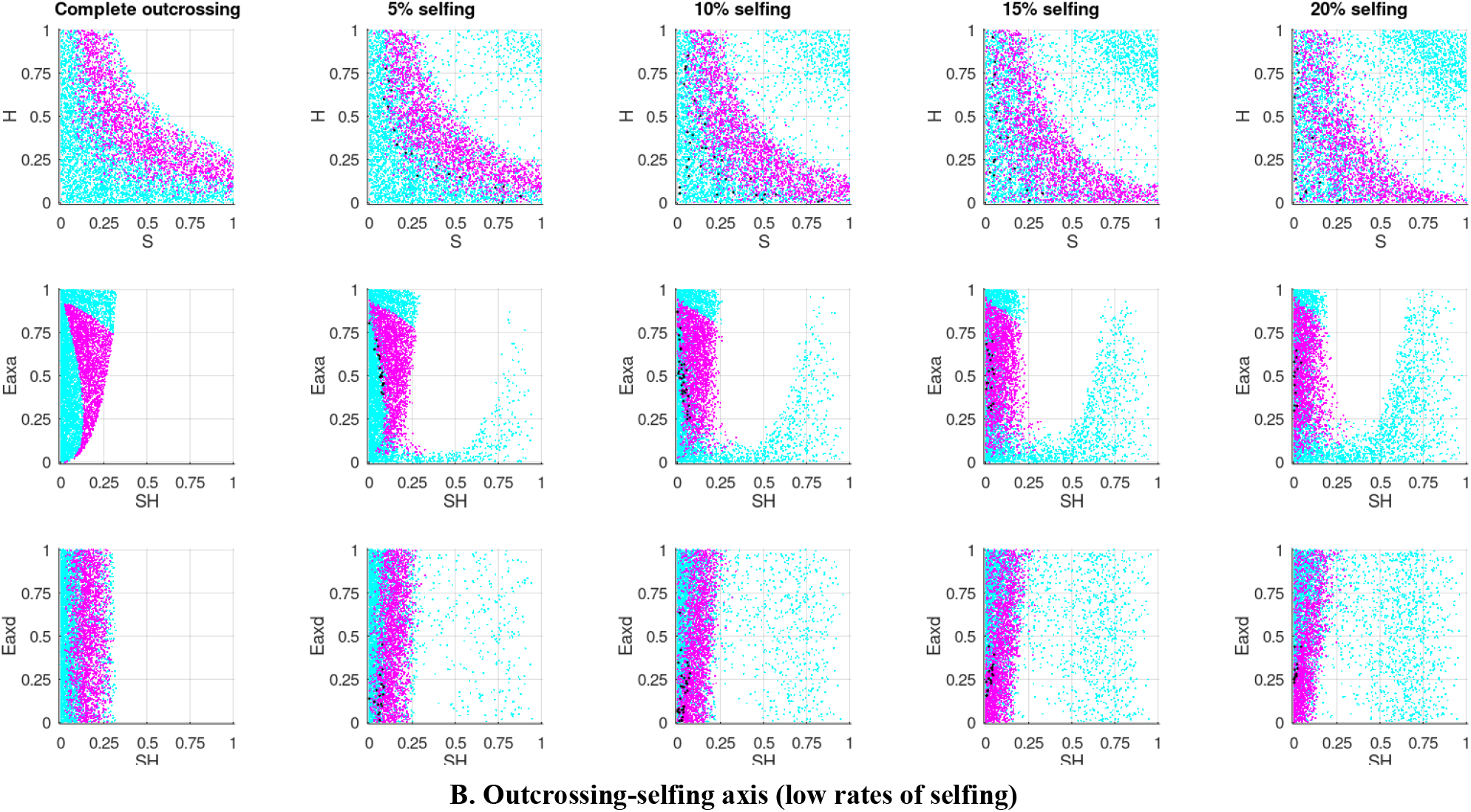
The effects of the main selection parameters on the evolutionary advantage/disadvantage of *fitness-dependent recombination* under different mating systems along the outcrossing–clonality (A) and outcrossing–selfing (B) axes. The rows of scatter plots within the axes represent the most informative coordinate plains: *S–H* (the first row), *SH–E_a×a_* (the second row), and (for the outcrossing-selfing axis only) *SH–E_a×d_* (the second row). The coloured points in the scatter plots represent only selection regimes that disfavour non-zero constant recombination (these correspond to the blue point in Fig. 1). Some of these regimes favour fitness-dependent recombination (magenta) while others disfavour it (cyan). Under low rates of selfing, there occur rare regimes that maintain a stable polymorphism between fitness-dependent recombination and zero recombination (black).

All these effects remained consistent along the *outcrossing–clonality axis*, with the decreasing values of *SH* compatible with selection for fitness-dependent recombination. In systems with partial selfing, the evolutionary advantage or disadvantage of fitness dependence under intermediate selfing was determined, first and foremost, by additive-by-dominance epistasis *E_a×d_*, selection against heterozygotes *SH* and triple product *SHE_a×d_*. The roles of additive-by-additive epistasis and the magnitude of recombination plasticity appeared secondary, although not negligible.

While usually one of the two competing modifier alleles (either for fitness-dependent recombination or zero recombination) moved towards fixation regardless of starting frequency, we occasionally observed more complicated outcomes. Situations with stable polymorphism occurred rarely (<1.2%) and only under relatively low (5–30%) rate of selfing. The corresponding parameter combinations were located along the border between the areas in which recombination was favoured and disfavoured (Fig. 3B, black points). We also observed sporadic regimes with bistability, where each of the two competing modifier alleles spread only when abundant.

In the area of realistic selective coefficients, the pattern was in general consistent with that observed over the full parameter space. Along the outcrossing–selfing axis, fitness-dependent recombination was favoured under relatively strong selection against heterozygotes *SH*, and this requirement relaxed with clonality (Fig. S2A). In systems with partial selfing, the evolutionary advantage/disadvantage of fitness-dependent recombination was strongly affected by both the additive-by-additive *E_a×a_* and additive-by-dominance *E_a×d_* epistatic components. However, in the area of realistic selection fitness-dependent recombination was favoured the most under relatively high rate of selfing, ~75–80% (Fig. S2B), while over the full parameter space this happened under much lower rate, ~15%.

### (c) Fitness-dependent recombination is often favoured even when constant recombination is not

To quantify how frequently fitness dependence preserves selection for recombination, we defined the ‘recombination-rescuing potential’ as the fraction of selection regimes favouring fitness-dependent recombination when non-zero constant recombination is disfavoured. Overall, the recombination-rescuing potential of fitness dependence under complete outcrossing was substantial: ~32–39% depending on the magnitude of recombination plasticity (Fig. 2A, green curve). Increasing clonality tended both to favour positive constant recombination and to increase the recombination-rescuing potential of fitness dependence, up to ~81–89% (Fig. 2A, blue and green curves). In contrast, along the *outcrossing–selfing axis*, the recombination-rescuing potential behaved non-monotonously: it first increased, reaching the maximal values of ~36–39% under ~15% selfing, before decreasing almost to zero under high rates of selfing (Fig. 2B, green curve).

After examining the whole parameter space, we focused on more realistic regimes. Along the *outcrossing–clonality axis*, the dynamics of the recombination-rescuing potential in these regimes were qualitatively similar to that found in the full parameter range (Fig. 2C, green curve). Along the *outcrossing–selfing axis*, the pattern was different: the recombination-rescuing potential was much more pronounced (up to >88%) under intermediate rates of selfing (20-80%) and declined under either low or high rates of selfing (Fig. 2D, green curve).

For both the entire parameter space and the area of realistic selection, we also considered systems with four and five selected loci. The recombination-rescuing potential of fitness dependence increased with the number of loci, although not drastically. Among systems with partial selfing, the effect of the number of loci was more pronounced under low-to-intermediate rates of selfing (Fig. S3).

### (d) The effect of fitness-dependent recombination on population-level parameters

We also examined the effect of fitness-dependent recombination on several population-level characteristics: (*i*) the time needed for the population to reach the equilibrium (state of mutation–selection balance), (*ii*) mean fitness in the equilibrium population, and (*iii*) standard deviation of fitness in the equilibrium population. No effect was found on the time needed to reach the equilibrium and a marginal effect was observed on the mean fitness and its variance (Fig. S4). This is consistent with the analogous results of other studies [8,13] and is expected given that the impact of fitness dependence is limited by the frequency of individuals carrying multiple mutations.

Regardless of whether fitness-dependent recombination was favoured or disfavoured, it typically ensured a higher mean fitness than zero recombination. However, under complete outcrossing, this increase in mean fitness was much larger when fitness-dependent recombination was disfavoured than favoured (~8·10^-10^ against ~2·10^-12^). This difference tended to diminish along the outcrossing–clonality axis. It also diminished along the outcrossing–selfing axis and even reversed: when the rate of selfing was >25%, fitness-dependent recombination increased mean fitness by more when it was favoured.

Under complete outcrossing, fitness dependence also led to a higher standard deviation of fitness compared with zero recombination. Again, this happened regardless of which of the two strategies was favoured, but when fitness-dependent recombination was disfavoured it increased the standard deviation more substantially (~10^-8^ against ~10^-11^). With a higher rate of selfing, fitness-dependent recombination increased the standard deviation when disfavoured and decreased it when favoured.

## 4. Discussion

Our simulations show that fitness-dependent recombination is often favoured under selection regimes that disfavour any non-zero constant recombination. This happens across a range of mating strategies, therefore extending our recent result [13] beyond panmixia. Moreover, together with similar earlier results on cyclical selection [12], these findings justify viewing fitness dependence as a potential generic factor increasing selection for recombination. The revealed recombination-rescuing potential of fitness dependence varies along the mating gradients, being more pronounced under higher rates of clonality and intermediate rates of selfing, especially under realistic values of selective coefficients. This variation probably ‘inherits’, at least partially, some more general regularities, namely those featured by the interplay between recombination *per se* (regardless of its possible dependence on genotype fitness) and mating strategy. However, to best of our knowledge, no solid theory was developed for this intriguing interplay except for a handful of theoretical simulations which have focused on the effect of selfing and clonality on the evolution of recombination [15–17]. We can therefore only provide speculative explanations for the above-presented results.

Under complete outcrossing, non-zero constant recombination appeared to be favoured when selection against heterozygotes was sufficiently strong (high *SH* values) (Fig. 1). In other words, non-zero constant recombination is selected for when mutations are considerably harmful (i.e., strongly deleterious and/or highly dominant). Given the negative epistasis assumed in the model, this can be explained within the mutational deterministic hypothesis: gathering several harmful mutations in one haplotype block indeed allows purging them more efficiently [18–20]. In such situations, recombination would increase the linkage disequilibrium between the selected loci by inflating the frequencies of homozygous multilocus haplotypes (i.e., those with either none or several mutations) at the expense of heterozygous ones (i.e., those with a single mutation). In turn, the increased linkage disequilibrium further ‘feeds’ purifying selection (see [21] for a review). This pattern became more pronounced with higher rates of clonality, so that non-zero constant recombination was favoured under smaller values of *SH* (Fig. 1A). Thus, selection against moderately harmful mutations that do not favour non-zero constant recombination under complete outcrossing, will favour it under high clonality. This suggests that asexuality ‘aggravates’ the deleterious effect of mutations, which largely agrees with Balloix et al. [22] who emphasized the harmful effect of deleterious mutations in highly-clonal organisms. Notably, the discrimination between regimes favouring/disfavouring fitness-dependent recombination followed the analogous pattern for non-zero constant recombination. Namely, modifiers conferring non-zero recombination (either constant or fitness-dependent) appeared to be favoured when the mutations were more deleterious and/or more dominant (Fig. 1A and 3A).

Intriguingly, complete selfing led to the opposite pattern: non-zero constant recombination was favoured when selection against heterozygotes was relatively week (low *SH* values) (Fig. 1B). Presumably, this happens due to extremely low heterozygosity. Selfing exponentially increases the rate of homozygosity and the fate of residual heterozygotes is defined by the interplay between selection intensity and mutation rate. Arguably, under complete selfing, only slightly harmful mutations (i.e., with low deleterious effect and/or low dominance) are retained in the population at frequencies high enough to induce selection on recombination modifiers, while more harmful mutations are quickly purged. In our simulations, intermediate rates of selfing favoured recombination (both constant and fitness-dependent) the most. This finding echoes the results obtained by Charlesworth et al. [15], and Holsinger and Feldman [16] arguing that partial selfing allows selection for non-zero constant recombination given certain viability matrices, in contrast to both complete outcrossing or complete selfing where such recombination-favouring regimes were not found [23,24]. Unfortunately, even the relatively simple setup (two-locus selected system with polymorphism maintained due to a heterozygote advantage) used in the above-mentioned models [15,16,23,24] appeared too complex to reveal in detail the effects of different parameters on the evolution of recombination. As Holsinger and Feldman inferred, “both the intensity of selection on modifiers of recombination and its direction depend in a complicated way not only on the degree of selfing but also on the viability system and other parameters”. Later, Roze and Lenormand [17] examined the evolution of recombination under mutation–selection balance. They showed that selfing, especially in the sporophytic form, strengthens the relative role of dominance-by-dominance epistasis up to unusual situation when recombination becomes favourable even under positive additive-by-additive epistasis. In our simulations, we did not observe significant changes in the relative role of dominance-by-dominance epistasis along the outcrossing–selfing axis - perhaps, since we overlooked the whole area of positive values of epistatic components. However, the role of additive-by-dominance epistasis did increase, and under intermediate rates of selfing it became more influential than additive-by-additive one (Table S1B).

It seems hard to infer to what extent the herein obtained results are consistent with empirical data. Several genome-wide comparisons of two *Arabidopsis* species have revealed on average higher recombination rate per physical length in the selfer (*A. thaliana)* than in the outcrosser (*A. lyrata)* [25–27]. Potentially, these estimates could be biased since genome size itself is known to be affected by the mating system, being on average lower in selfers [28]. Nevertheless, more cautious (treating genome size as a covariate) and more extensive (covering 142 angiosperm species) analysis performed by Ross-Ibarra has confirmed that selfing generally favours recombination [29]. Yet, understanding the selfing-recombination interplay may be much more challenging than just controlling for genome size. The mating system can affect not only recombination but also other variation-related traits, such as mutation rate and the activity of transposable elements [28]. Despite some insights into the coevolution of recombination and these traits (for example, recombination rate usually negatively correlates with the activity of transposable elements [30]), it remains poorly understood. In our simulations, selection regimes favouring non-zero constant recombination under selfing, but disfavouring it under outcrossing, emerged when the mutations were slightly-to-moderately harmful (Fig. 1B). It seems plausible that most of the mutations in nature are exactly of this kind. If so, then our theoretical results are indeed consistent with the empirically established a higher recombination rate in selfers.

## Acknowledgements

The authors express gratitude to the Hypernet Labs team, and especially Jennifer Hudson, for the Galileo platform for remote computations (https://hypernetlabs.io) and their kind support. SR is deeply thankful to Nataliya Rybnikova for her assistance in surveying the literature.

