## Supplementary material for "Fitness dependence preserves selection for recombination across diverse mixed mating systems": S1-S4: S1.docx

**Table S1A. The effects of the selection parameters on the evolutionary advantage/disadvantage of *non-zero constant recombination* under mating systems alone the *outcrossing–clonality axis* (based on binary logistic regression with no factor interactions)**

| Selection  parameter | Complete outcrossing | | 25% clonality | | 50% clonality | | 75% clonality | | 99% clonality | |
| --- | --- | --- | --- | --- | --- | --- | --- | --- | --- | --- |
|  | B | W | B | W | B | W | B | W | B | W |
| Intercept | 14.123 | 1374.922 | 12.390 | 1389.897 | 9.580 | 1382.954 | 6.405 | 1098.406 | 1.231 | 25.504 |
| *S* | -15.514 | 1521.437 | -14.659 | 1612.832 | -12.451 | 1776.433 | -9.819 | 1763.293 | -8.721 | 302.386 |
| *H* | -15.686 | 1521.318 | -14.728 | 1605.057 | -12.404 | 1765.844 | -9.829 | 1734.863 | -7.572 | 305.804 |
| *E_a×a_* | 5.634 | 780.668 | 4.840 | 701.913 | 3.704 | 569.646 | 2.053 | 235.737 | 0.235 | 1.035 |
| *E_a×d_* | 0.127 | 0.719 | -0.007 | 0.002 | -0.228 | 2.879 | -0.225 | 3.138 | -0.054 | 0.055 |
| *E_d×d_* | -0.007 | 0.002 | 0.173 | 1.433 | 0.171 | 1.608 | 0.020 | 0.024 | -0.209 | 0.820 |
| AIC | 3505 | | 3770 | | 4313 | | 4769 | | 1597 | |
| Accuracy, % | 92.1 | | 91.4 | | 89.7 | | 88.8 | | 97.9 | |
| AUC | 0.981 | | 0.978 | | 0.971 | | 0.956 | | 0.941 | |

**Table S1B. The effects of the selection parameters on the evolutionary advantage/disadvantage of *non-zero constant recombination* under mating systems alone the *outcrossing–selfing axis* (based on binary logistic regression with no factor interactions)**

| Selection  parameter | Complete outcrossing | | 25% selfing | | 50% selfing | | 75% selfing | | Complete selfing | |
| --- | --- | --- | --- | --- | --- | --- | --- | --- | --- | --- |
|  | B | W | B | W | B | W | B | W | B | W |
| Intercept | 14.12 | 1374.92 | 2.44 | 609.72 | -0.457 | 22.133 | -3.693 | 883.663 | -11.575 | 1448.395 |
| *S* | -15.51 | 1521.44 | -3.95 | 1733.6 | -0.852 | 92.379 | 3.192 | 849.576 | 12.934 | 1658.933 |
| *H* | -15.69 | 1521.32 | -0.25 | 9.057 | 2.675 | 796.205 | 5.412 | 1817.143 | 13.285 | 1679.89 |
| *E_a×a_* | 5.63 | 780.67 | -2.58 | 848.969 | -4.477 | 1804.352 | -4.932 | 1586.475 | -5.898 | 949.798 |
| *E_a×d_* | 0.13 | 0.72 | 1.55 | 331.489 | 1.815 | 397.869 | 1.519 | 228.126 | 0.085 | 0.372 |
| *E_d×d_* | -0.007 | 0.002 | -0.10 | 1.503 | 0.011 | 0.016 | 0.058 | 0.345 | -0.005 | 0.001 |
| AIC | 3505 | | 10566 | | 9573 | | 7845 | | 4031 | |
| Accuracy, % | 92.1 | | 74.3 | | 76.9 | | 83.6 | | 92.1 | |
| AUC | 0.981 | | 0.811 | | 0.829 | | 0.894 | | 0.976 | |

Both tables represent selection regimes with all five parameters (*S*, *H*, and three epistasis coefficients) simulated as evenly distributed in the interval (0, 1).
