## Supplementary material for "Fitness dependence preserves selection for recombination across diverse mixed mating systems": S1-S4: S2.docx

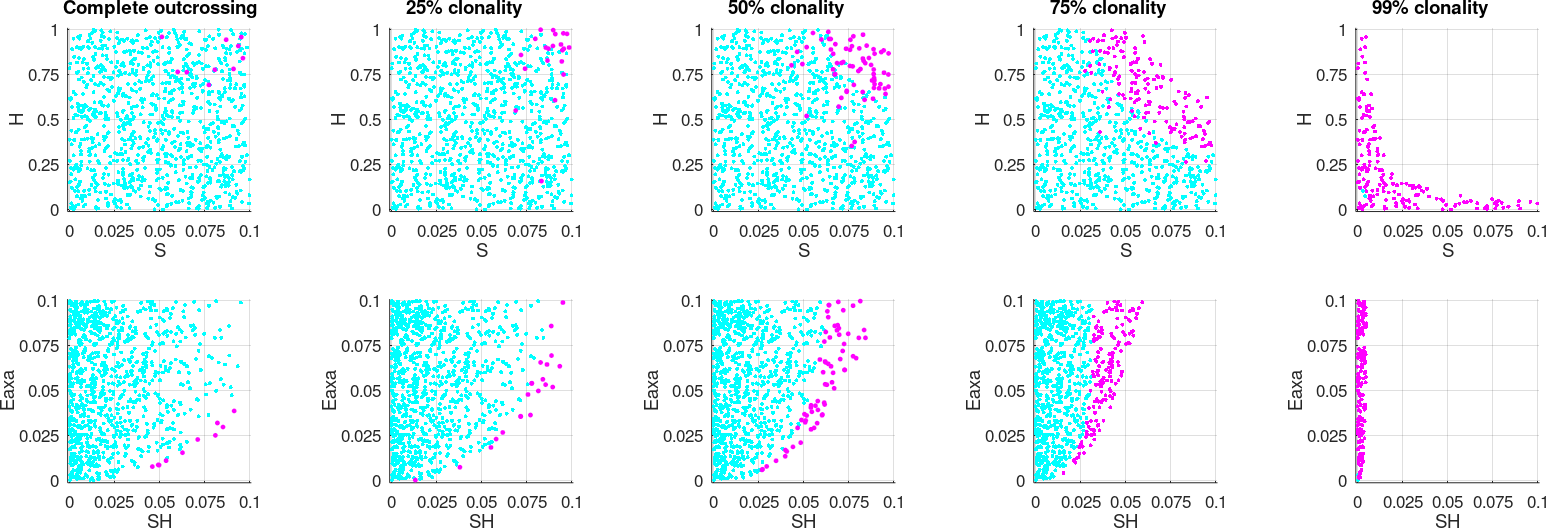

**A. Outcrossing-clonality axis**

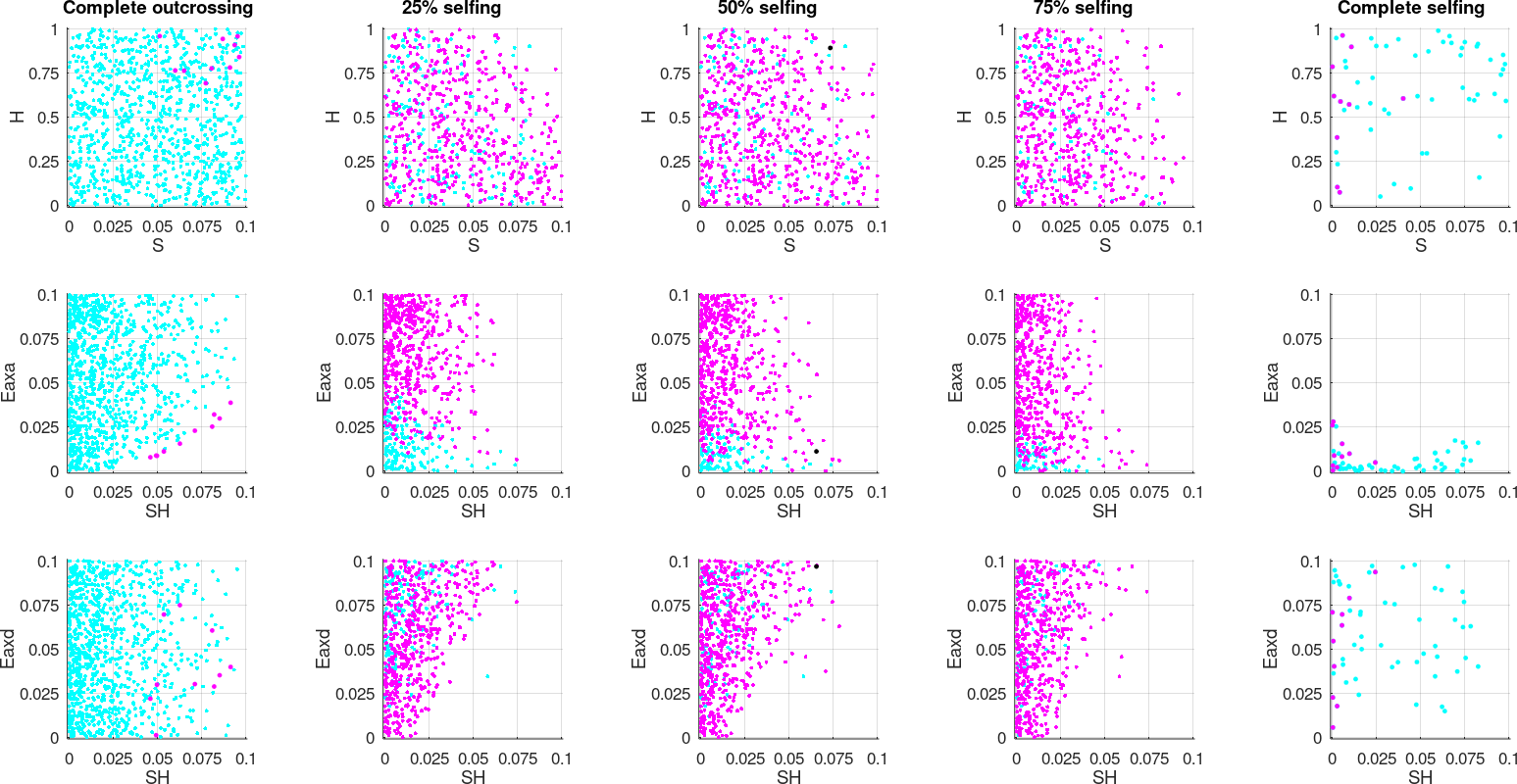

**B. Outcrossing-selfing axis**

**Figure S2: The effects of the main selection parameters *within biologically realistic ranges* on the evolutionary advantage/disadvantage of *fitness-dependent recombination* under different mating systems along the outcrossing–clonality (A) and outcrossing–selfing (B) axes.** The rows of scatter plots within the axes represent the most informative coordinate plains: *S–H* (the first row), *SH–E_a×a_* (the second row), and (for the outcrossing–selfing axis only) *SH–E_a×d_* (the second row). The coloured points in the scatter plots represent only selection regimes that disfavour non-zero constant recombination. Some of these regimes favour fitness-dependent recombination (magenta) while others disfavour it (cyan).
