## Supplementary material for "Fitness dependence preserves selection for recombination across diverse mixed mating systems": S1-S4: S3.docx

| 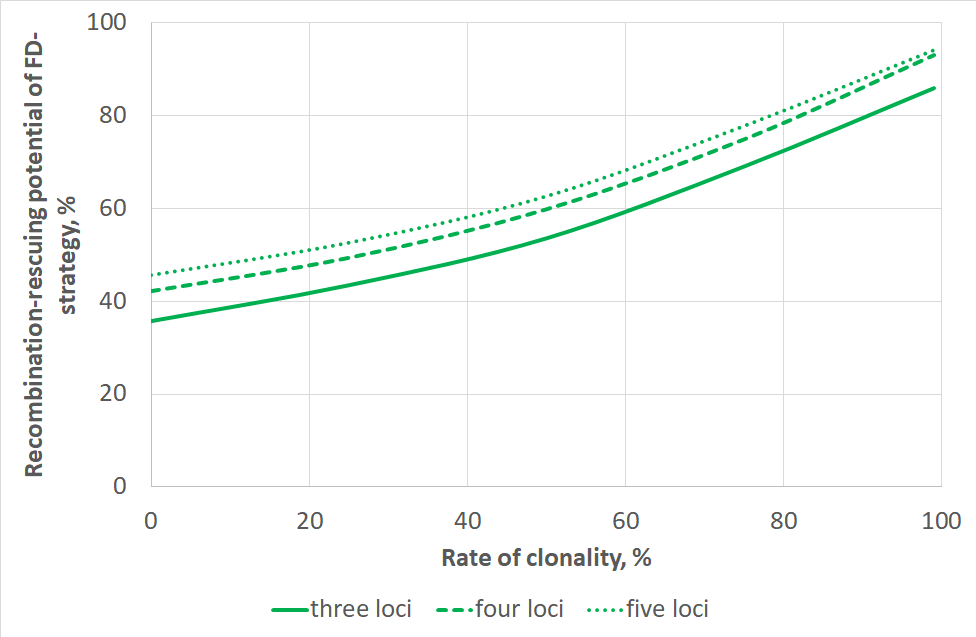 | 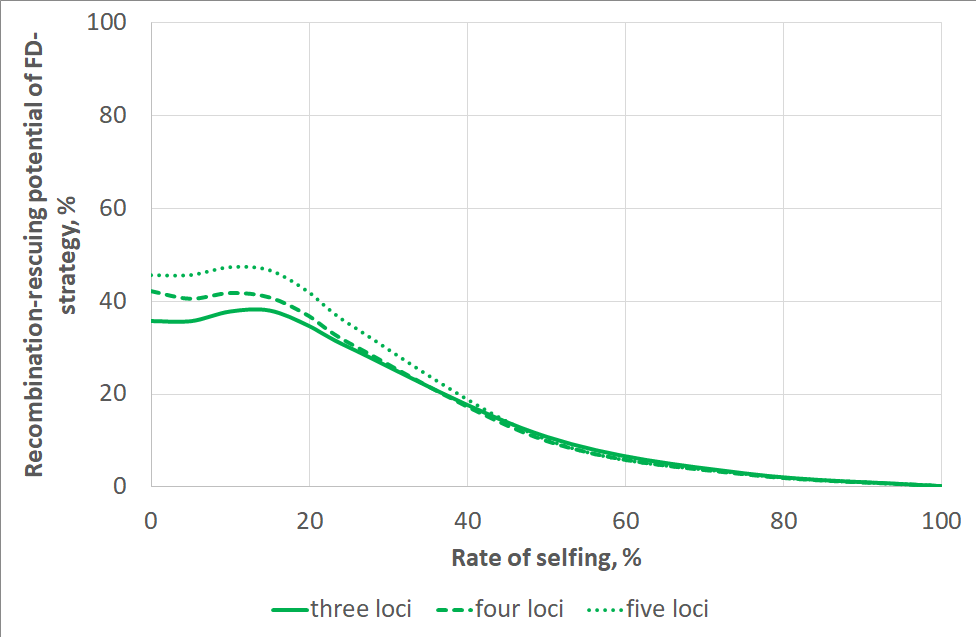 |
| --- | --- |
| **A. Outcrossing-clonality axis (all selection regimes)** | **B. Outcrossing–selfing axis (all selection regimes)** |
| 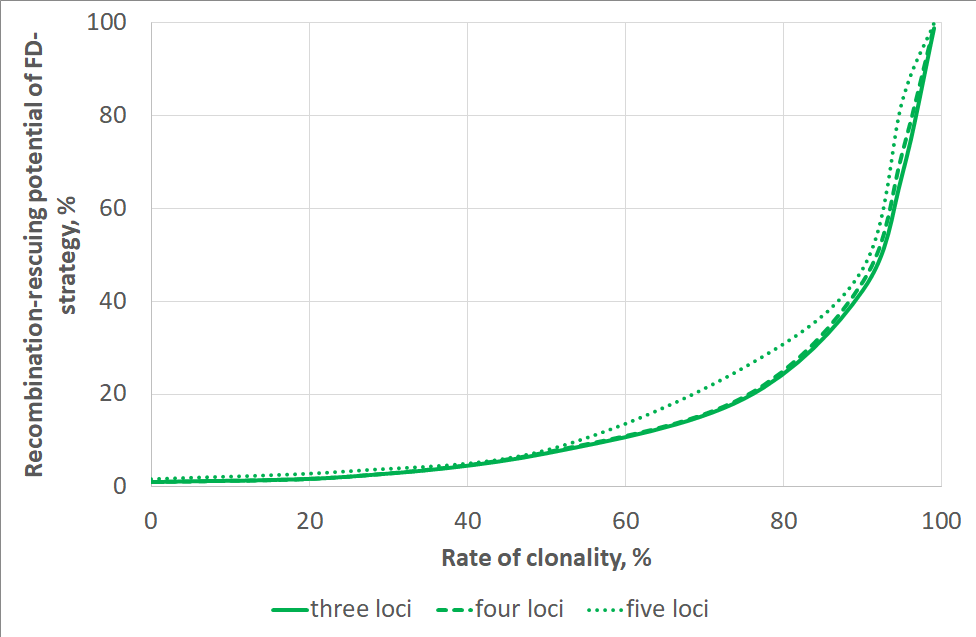 | 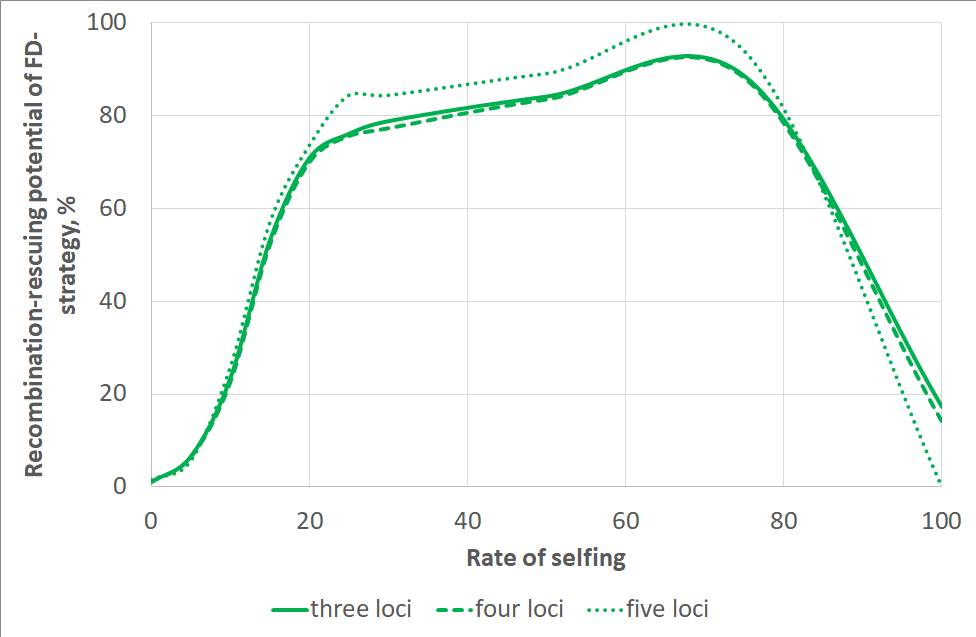 |
| **C. Outcrossing-clonality axis (weak-to-moderate selection regimes)** | **D. Outcrossing–selfing axis (weak-to-moderate selection regimes)** |

**Figure S3: The effect of the number of selected loci on the recombination-rescuing potential of fitness-dependent recombination, mating systems along the outcrossing–clonality (A and C) and outcrossing–selfing (B and D) axes**
