## Supplementary material for "Fitness dependence preserves selection for recombination across diverse mixed mating systems": S1-S4: S4.docx

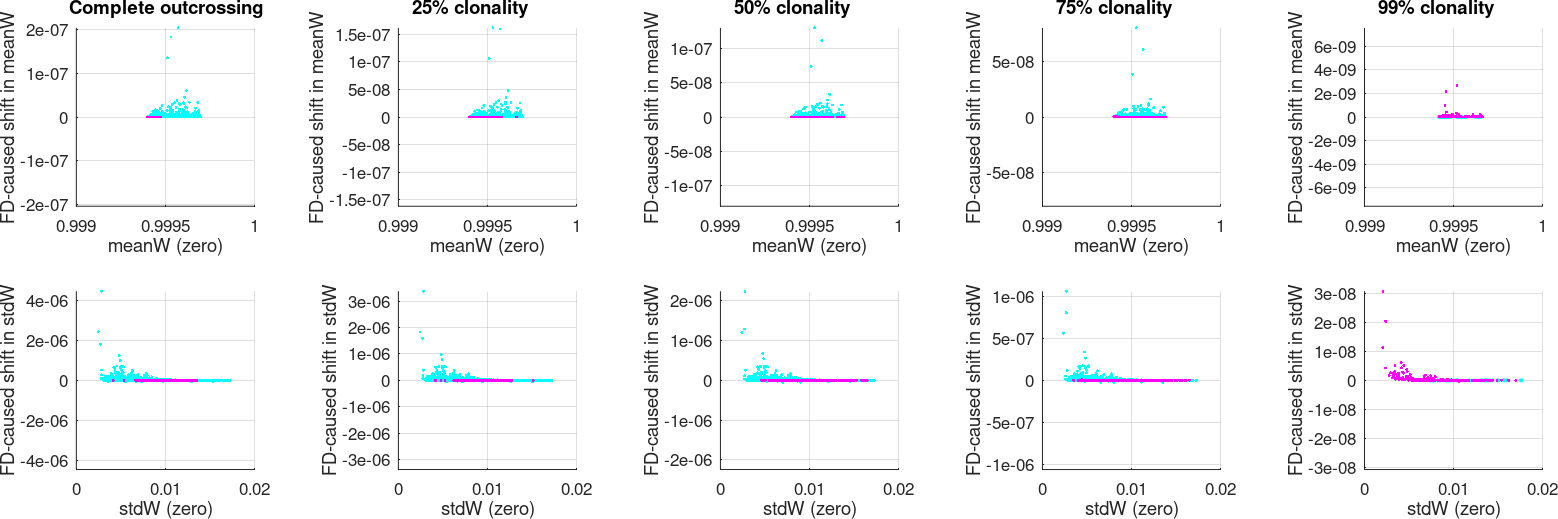

**A. Outcrossing-clonality axis (all selection regimes)**

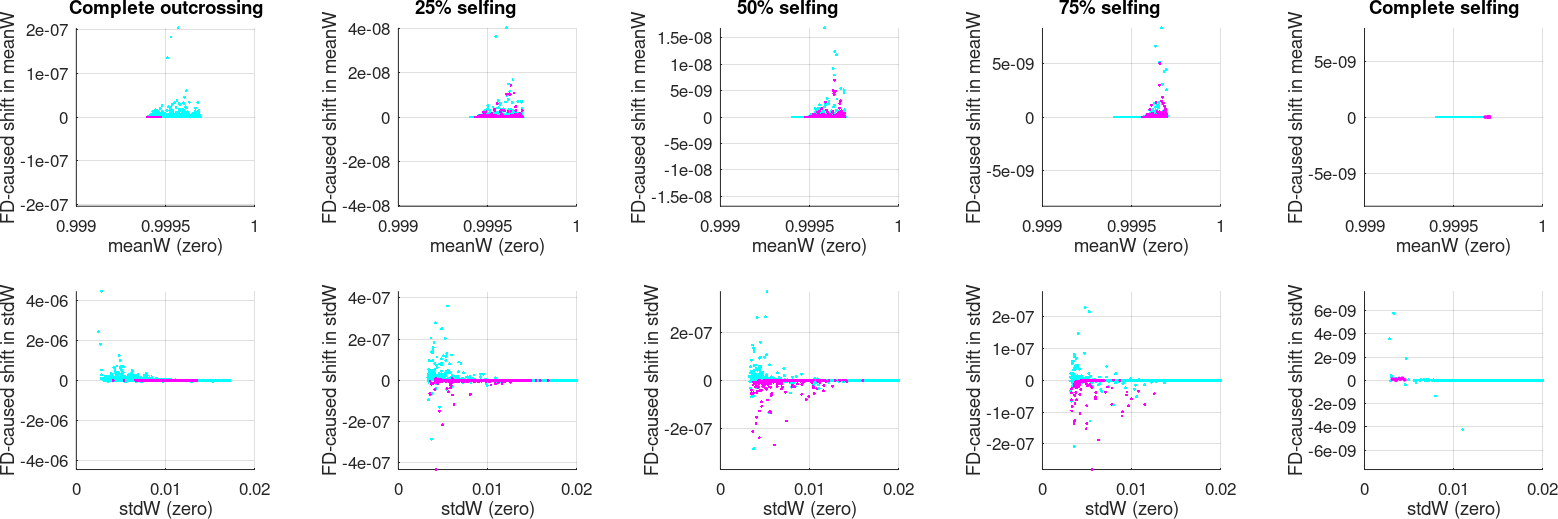

**B. Outcrossing-selfing axis (all selection regimes)**

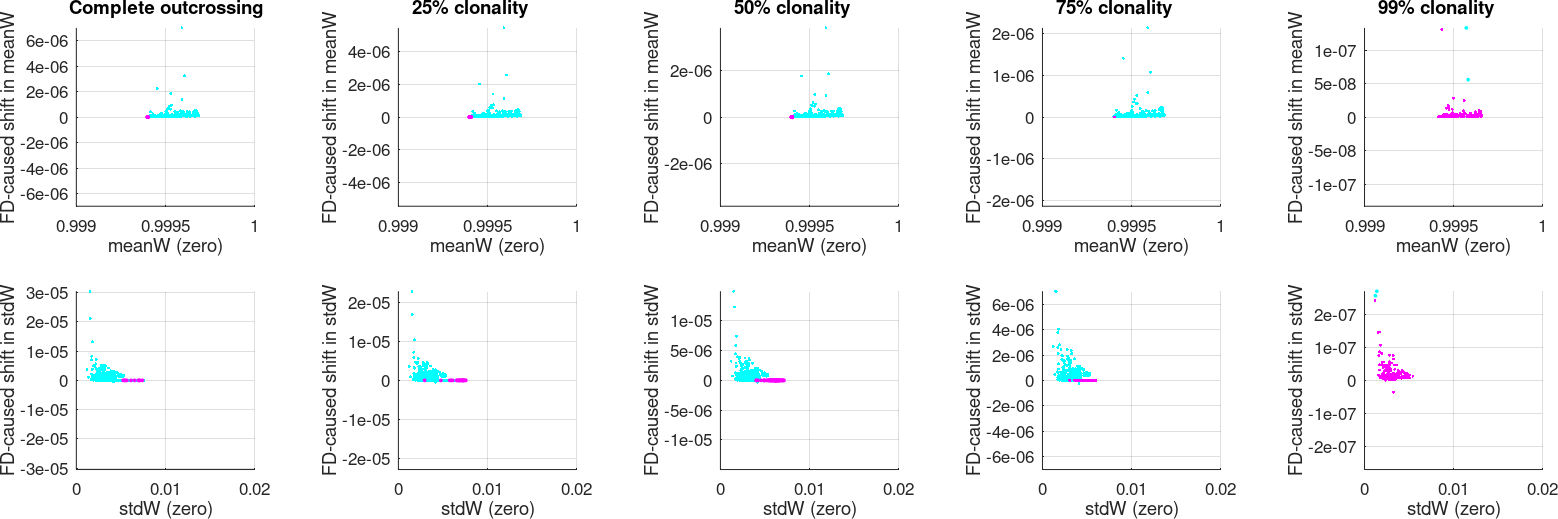

**C. Outcrossing-clonality axis (selection regimes with *S* and *E*<0.1)**

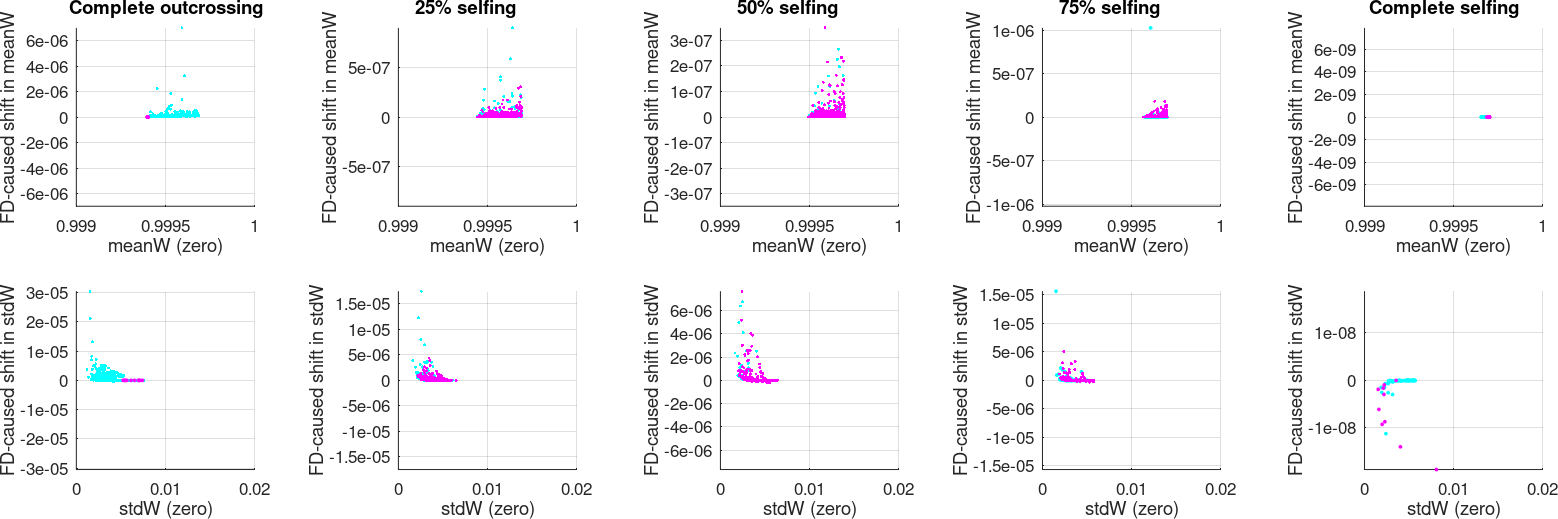

**D. Outcrossing-selfing axis (selection regimes with *S* and *E*<0.1)**

**Figure S4: The effects of fitness dependence on population-level characteristics under different mating systems along the outcrossing–clonality (A and C) and outcrossing–selfing (B and D) axes.** The rows in each subplot represent mean (the top row) and standard deviation (the bottom row) of fitness in the population. In each scatter plot, the *x*-axis stands for the value under zero recombination while the *y*-axis shows the shift caused by fitness dependence. The coloured points represent only selection regimes that disfavour non-zero constant recombination. Some of these regimes favour fitness-dependent recombination (magenta) while others disfavour it (cyan).
